# Sensing endogenous RNA in living human cells using a CRISPR-activated protease

**DOI:** 10.1101/2024.07.17.603943

**Authors:** Katja Blumenstock, Alexander Hoch, Leo D. Hinterlang, Caroline I. Fandrey, Niels Schneberger, Gregor Hagelueken, Jonathan L. Schmid-Burgk

## Abstract

Most techniques used to detect specific mRNAs in eukaryotic cells require to extract nucleic acids and thereby kill the cells. A programmable sensor for monitoring endogenous transcripts in living cells, in contrast, would enable to enrich living cells based on a specific transcription or splicing event, and studying these cells by live microscopy or sequencing methods requiring intact cells. We have engineered CRISPR-READ, a live cell RNA detector based on the CRISPR-associated Lon protease CalpL and a cA_4_-producing Type III CRISPR system. Upon RNA-programmable RNA sensing, CRISPR-READ produces an orthogonal second messenger, which leads to the cleavage of a dual FRET / localization reporter compatible with FACS sorting and live microscopy. Using this genetically encoded sensing circuit as a readout for a genome-wide CRISPR perturbation screen, we identified an extended Type-I interferon signaling cascade; RNA-Seq on sensor-sorted cells enabled unbiased identification of correlated stochasticity in gene expression across single cells.

## INTRODUCTION

Comprehensively mapping transcriptional signaling networks and thereby identifying key signaling hubs will enable to elucidate etiologies of genetic diseases and to identify novel drug targets. Therefore, methods for measuring gene expression changes and correlating them with cellular functions as well as perturbations are essential for mapping biological *wiring diagrams* determining state transitions and stochastic processes in human cells. Despite the critical role of monitoring transcript dynamics in combination with other live-cell readouts, available techniques require - with very limited exceptions discussed below - to kill the cells: While sequencing-^1 2^ or hybridization-based (HCR-FISH ^3 4^, flow-FISH ^5 6 7^) gene expression measurements require to obtain access to the RNA inside of the cells, affinity reagents can monitor gene expression at the protein level only on the cell surface without significantly disrupting cellular integrity. Engineered RNA editing-based sensors like RADARS ^8 9 10^ offer a snapshot of sequence-specific RNA expression levels in living cells, but their limited sensitivity and complex design constraints precluded application to single-cell measurements of meaningful biological state changes and correlating them with cellular function so far. Lastly, genetic reporters coupling transgene expression to an endogenous promoter are highly laborious to create, and do not recapitulate the full life cycle of an RNA transcript without interfering with it; furthermore, these reporters inherently convolve RNA abundance information with protein life cycle effects. This lack of technologies largely precluded correlating the abundance of specific RNA sequences with single live cell functions, or high-quality full-length RNA sequencing of cells enriched for a specific transcriptional state. To develop a truly programmable and sensitive RNA-specific readout in single live human cells, we therefore turned to natural biological systems that perform a similar task, recognizing defined nucleic acid sequences and executing a signaling program in bacterial cells without killing them.

CRISPR immune systems appeared to be an ideal starting point for this endeavor. They are adaptive bacterial immune systems that allow bacteria to defend themselves against mobile genetic elements (MGEs), such as phages and plasmids. In a nutshell, the bacterial Cas1-Cas2 complex captures and incorporates small snippets of invader DNA into a chromosomal CRISPR locus that acts as a memory of previous invasions. Transcripts of this locus are processed into small CRISPR RNAs (crRNA), which are then integrated into large ribonucleoprotein recognition complexes such as the well-known Cas9, which allow the organism to detect and destroy MGEs ^11 12^. Unlike type I and II CRISPR systems, which degrade invader DNA directly, the recognition complex of type III systems recognizes transcription products of foreign DNA. One component of this complex, Cas10, produces unique cyclic oligonucleotide (cOA) second messengers to activate various downstream effector proteins with cOA sensor domains ^13 14^. The effector proteins have diverse functions, including DNAses, RNAses, transcription regulators and, as very recently discovered, proteases ^15 16 17 18 19^. The dependence on a second messenger is an elegant way to activate complex cellular machineries in a fast, diffusion-limited way and to amplify the signal. It is therefore not surprising that cyclic nucleotides are common in immune signaling in both eukaryotes and prokaryotes. For instance, the human cGAS-STING pathway uses cyclic 2’,3’-GMP-AMP (cGAMP) to control antiviral immunity ^20 21 22^. Other prominent examples are cAMP and cGMP. In addition to CRISPR, bacteria use much simpler systems, like CBASS, to combat phage attacks ^23 24 25^. We assumed that the CRISPR controlled CalpL protease from the thermophilic *Sulfurihydrogenibium* sp. could be an ideal candidate for our envisioned sensor design ^19^. It is a small and very stable (60 kDa) Lon protease^26 27^ that is activated by cyclic tetraadenylate (cA_4_), which binds to its SAVED domain. This leads to oligomerization of CalpL and activation of the protease activity. Once activated, CalpL cleaves the anti-sigma factor CalpT, a small 33 kDa protein, into two defined peptides. Notably, CalpL and CalpT form a stable complex, which is needed for protease activity.

We have used the unique properties of the CalpL / T complex to develop **CRISPR-READ** (<u>CRIS</u>PR-mediated <u>P</u>rogrammable <u>R</u>NA <u>R</u>eadout <u>E</u>nabled by cyclic <u>A</u>denylate <u>D</u>etection), a genetically encoded, cA_4_ activated and programmable RNA sensor, which leads to the cleavage of a dual FRET / localization reporter compatible with FACS sorting and live microscopy. We demonstrate the usability and power of the system on a number of challenging test cases.

## RESULTS

### Sequence-specific RNA sensing in living cells

To install an orthogonal signaling pathway in human cells which is activated in a sequence specific manner by cytosolic RNA, we combined two well-characterized Type III CRISPR systems from different prokaryotic species: Upon target recognition of a CRISPR sensing complex from *Thermus thermophilus sp. HB8*, ATP is converted to cA_4_, which binds to and activates a CRISPR-activated Lon protease from *Sulfurihydrogenibium* sp. YO3AOP1 (Fig. 1A). We chose this design as we reasoned that cA_4_ has no known function in human cells, so it would spread the sensed signal throughout the cell without interfering with cellular signaling. To enable flexible readouts, we designed a fluorescent reporter consisting of mCherry, CalpT, and EGFP (Fig. 1B), inspired by proven TEV protease reporter designs ^28^. Proteolytic cleavage of this reporter can be detected by loss of FRET between two fluorescent proteins as well as by subcellular localization upon separation (Fig. 1C). To test the sequence-specific activation of this biological circuit, we first expressed matching and mismatching combinations of crRNAs and target RNAs from separate U6 promoters, along with CRISPR Cmr 2, 3, 4, 5 proteins. The target sequences, which were chosen without strong similarity to the human transcriptome, only activated the reporter when a reverse-complementary crRNA was expressed in the same cells (Fig. 1D), confirming a high specificity of CRISPR-mediating sensing ^29^.

**Figure 1.**
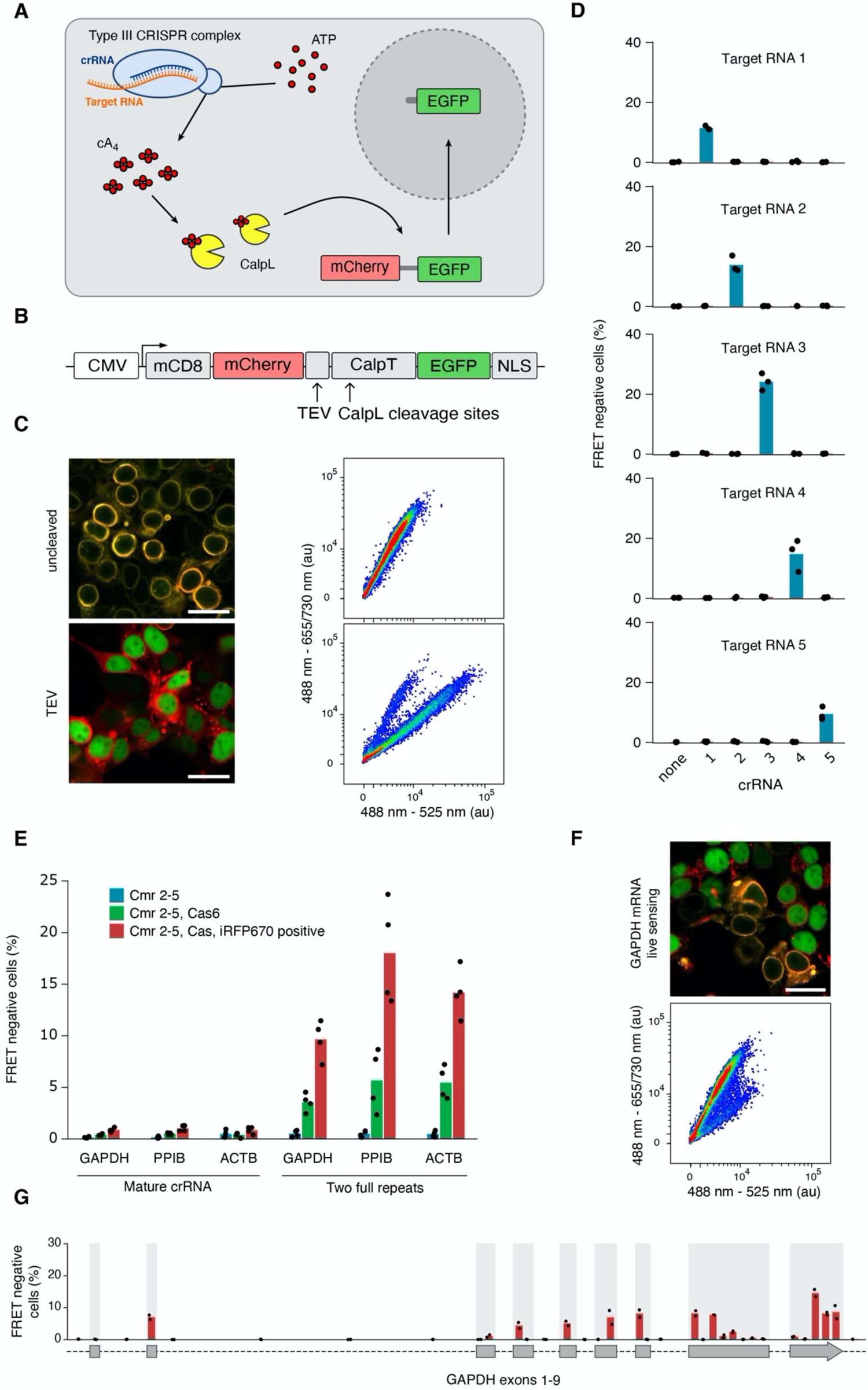
A combination of two Type III CRISPR systems can sense mRNAs in living human cells. **(A)** Schematic of programmable RNA sensing in human cells. Upon target hybridization of a Type III CRISPR complex from *Thermus thermophilus sp. HB8*, ATP is converted to cA_4_, which binds to and activates the CRISPR-activated Lon protease from *Sulfurihydrogenibium* sp. YO3AOP1. Subsequent proteolytic cleavage of a dual fluorescent reporter can be detected by visible light. **(B)** Schematic of the Lon protease reporter operating in living human cells. A CalpT domain from *Sulfurihydrogenibium* sp. YO3AOP1 as well as a TEV cleavage site for positive control experiments are inserted between red and green fluorescent proteins, which upon cleavage loose FRET activity. Furthermore, membrane- and nuclear localization domains separate the two colors upon cleavage. **(C)** Intracellular localization and FRET activity of the dual-color reporter in response to TEV-mediated intracellular cleavage in HEK 293T cells. Scale bar, 20 μm. **(D)** Sequence-specific programmable RNA sensing upon expression of matching and mismatching pairs of crRNAs and target RNAs under a polymerase III promoter. Shown are fractions of FRET-negative cells measured by FACS. **(E)** Optimization of CRISPR-READ for sensing endogenously expressed mRNAs. Reporter activation is shown across three crRNAs targeting endogenous genes, either as mature crRNAs or flanked by two full repeats, with or without Cas6, and with optional gating on iRFP670-expressing cells as a transfection control. **(F)** Intracellular localization and FRET activity of CRISPR-READ after sensing endogenously expressed GAPDH in HEK 293T cells. Scale bar, 20 μm. **(G)** Sensing efficiency of 34 crRNAs tiling the GAPDH pre-mRNA sequence; grey boxes indicate annotated exons. Additional tiling data are shown in Supplementary Fig. 1.

### Endogenous mRNA sensing in living cells

When we expressed crRNAs to sense three endogenous house-keeping mRNAs without over-expressing their cognate target sequences, however, no reporter activation could be detected (Fig. 1E, left panel blue bars). To enable sensing endogenous mRNAs, we reasoned that expressing mature crRNAs from a U6 promoter will always include at least one 5’ G and several 3’ U bases that do not match the natural configuration, thereby potentially limiting the efficiency of RNA sensing. To circumvent this problem, we expressed an immature crRNA with full CRISPR repeats on both sides of the target-specific spacer region, and co-expressed Cas6 protein for maturation. Strikingly, only the combination of full-length repeats and Cas6 expression led to reporter activation, sensing three endogenously expressed transcripts (Fig. 1E, right panel, green bars and Suppl. Fig. 1A). Additionally, gating cells for successful transfection by iRFP670 co-expression further increased the fraction of reporter-activated cells by more than 2-fold (Fig. 1E, red bars). Utilizing this optimized system, we found 9 out of 34 crRNAs targeting GAPDH pre-mRNA to lead to reporter activation in >5% of cells, while recapitulating the exon-intron structure of the mature mRNA, as all intronic crRNAs failed to activate the reporter. This observation is in line with the notion that mature spliced transcripts are sensed in the cytosol. A similar success rate was observed when sensing additional endogenous mRNAs (Suppl. Fig. 1B-C), with no strong sequence features predicting sensing efficiency (Suppl. Fig. 1D).

### Programmable CRISPR sensing of interferon stimulated genes (ISGs)

In order to test if the programmable CRISPR sensor can detect dynamic changes of transcript levels, we designed crRNAs to detect two well-studied ISGs as well as their reverse-complement sequences as controls. When stimulating the sensor-expressing cells with IFNβ, the fraction of reporter-activated cells was increased in a dose-dependent manner for both ISGs, while reverse-complement sequences were not sensed even at the highest concentration of IFNβ (Fig. 2A-B). Among target-expressing cells, CRISPR-READ was able to discern stochastic differences on single-cell level down to RPKM values of 100 (Fig. 2C). To study the genetic basis of dynamic transcriptome changes, we introduced a third CRISPR system from *Streptococcus pyogenes* to inactivate genes before ISG induction. First, we delivered two sgRNAs targeting the receptor for IFNβ or an unrelated receptor together with SpCas9 to generate permanent knock outs in a pool of cells. We then sorted the cells for transcriptome changes sensed upon IFNβ stimulation, and measured the genomic mutation rates at the targeted loci by Next-Generation Sequencing (NGS) (Fig. 3A). Indeed, genomic editing rates were highly skewed towards lower IFNβ receptor perturbation in reporter-activated cells, and higher editing in non-activated cells, while sensing GAPDH as an interferon independent control transcript did not skew knock out rates (Fig. 3B).

**Figure 2.**
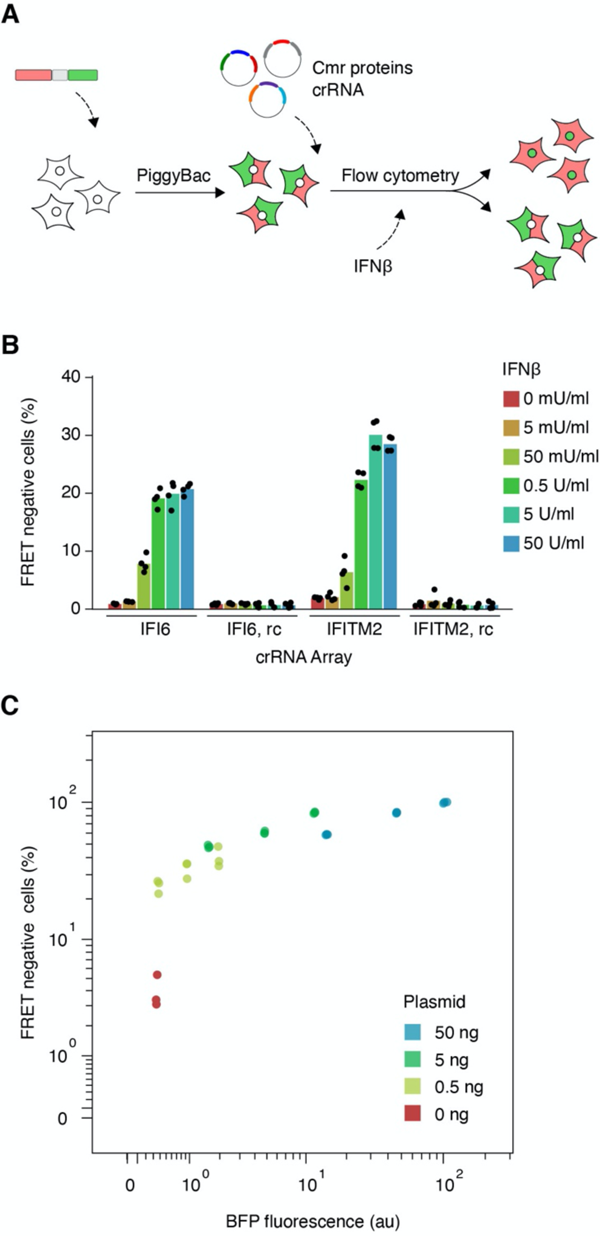
Detecting dynamic interferon receptor signaling in human cells using programmable CRISPR sensing. **(A)** Schematic of programmable sensing of Interferon Stimulated Gene (ISG) transcripts. **(B)** Titration of interferon beta on HEK 293T cells equipped with CRISPR -READ targeting indicated ISG mRNAs, or their reverse-complement sequences. Reporter activation was measured by flow cytometry. (**C**) Titration of a BFP-IFI6 expression plasmid transfected into HEK 293T cells equipped with CRISPR-READ targeting IFI6. Cells were analyzed by flow cytometry and were computationally split into three bins of BFP expression. Reporter activation rates are displayed in correlation to each bin’s mean BFP fluorescence. Mean RPKM expression levels across the bins were: red 1.44 ± 1.01; yellow 8.07 ± 2.55; green 158.65 ± 6.21; blue 1100.98 ± 66.35.

**Figure 3.**
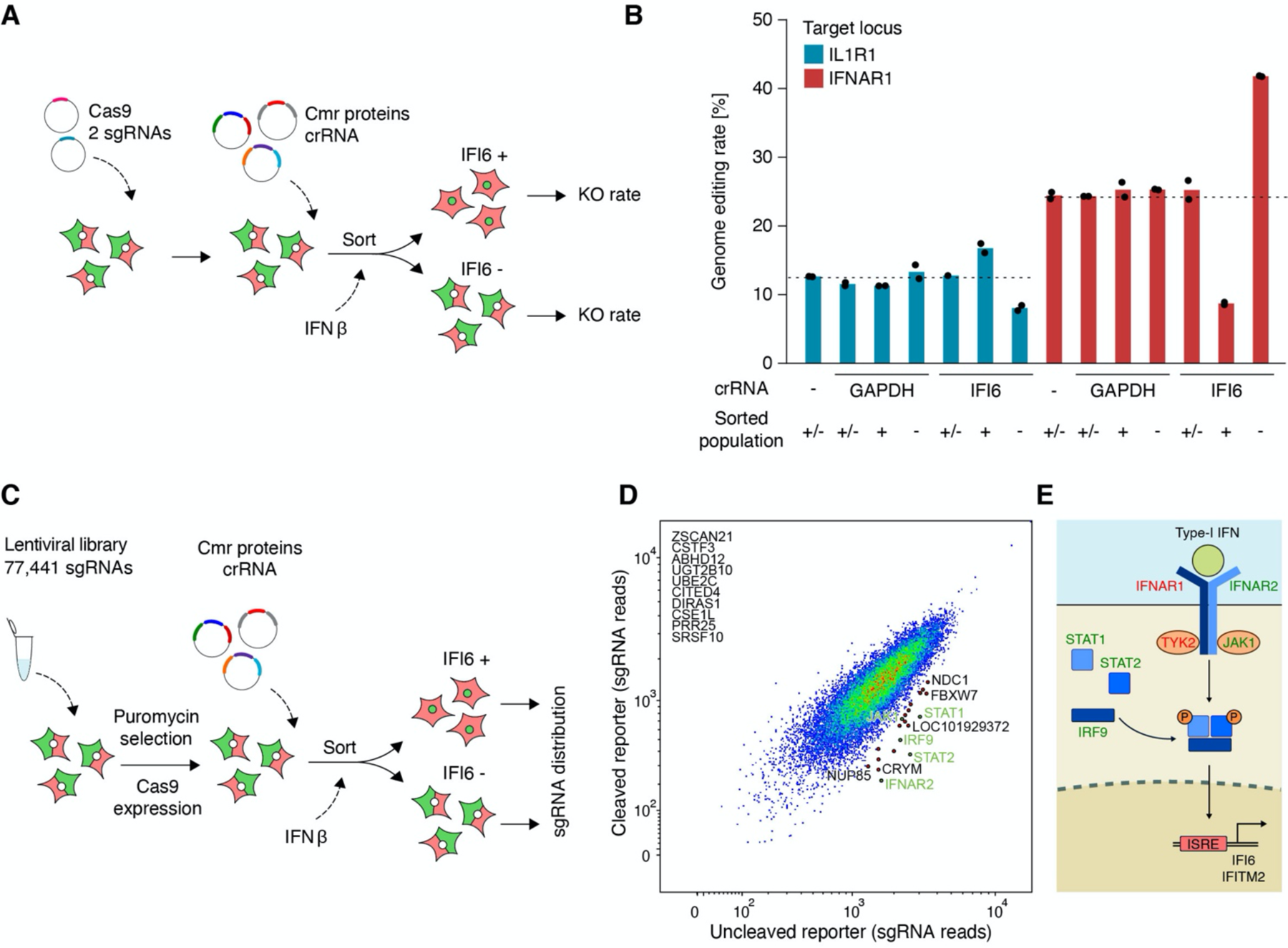
Genetic screening for signaling pathway members using programmable CRISPR sensing. **(A)** Outline of a pilot CRISPR perturbation screen based on programmable mRNA sensing as the readout. **(B)** Pilot perturbation screen consisting of two SpCas9 sgRNAs, targeting either the Type-I interferon receptor, or the unrelated IL-1 receptor. Upon interferon stimulation, cells were enriched based on programmable sensing of either a house-keeping gene (GAPDH), or an interferon-induced gene (IFI6). Shown are locus-specific genome editing rates in sorted populations as measured by NGS. **(C)** Outline of a genome-scale CRISPR perturbation screen based on programmable mRNA sensing as the readout. **(D)** Genome-scale perturbation screen combining CRISPR systems from three bacterial species; as outlined in (C), cells were perturbed in a pooled format using a lentiviral sgRNA library and transient SpCas9 expression. Upon interferon stimulation, programmable sensing of ISG-expression was used to enrich cells with a functioning or non-functioning IFN sensing pathway. The sgRNA representation was measured across sorted populations in eight replicates. Genes are colored based on their known function in relation to interferon-dependent signaling. **(E)** Pathway diagram of Type-I interferon receptor signaling, indicating known components identified by CRISPR-READ genome-scale perturbation screening.

### Genome-scale detection of co-regulated as well as regulating genes

Using programmable sensing of dynamic ISG induction as the readout, we next performed a genome-wide CRISPR screen to delineate pathway members regulating ISG induction. After introducing a library of 76,441 sgRNAs perturbing most human protein coding genes as well as 1,000 non-targeting control sgRNAs, cells were equipped with a CRISPR sensor, stimulated with IFNβ, and sorted for IFI6 mRNA sensing (Fig. 3C). The distribution of perturbing sgRNAs was highly skewed in IFI6 mRNA-negative versus positive cells towards genes known to be critically involved in the IFNβ sensing pathway, like IFNAR2, STAT1/2, JAK1, and IRF9 (Fig. 3D-E). Multiple additional hits were enriched that are involved in nuclear pore shuttling, which is crucial for transcription induction downstream of STAT1/2 phosphorylation.

The functional consequences and true randomness of noisy gene expression remain hard-to-study phenomena in biology due to the lack of single-cell measurements that do not create noise themselves. To directly identify co-regulated noisy genes across single cells, we programmed a CRISPR sensor to detect a transcript which we had previously observed to support reporter activation only in a fraction of cells upon low-level IFNβ stimulation (Fig. 2B). As CRISPR RNA sensing enables to retrieve intact cells for downstream analysis, we purified bulk cellular RNA from populations sorted for high or low sensing of IFI6 or GAPDH mRNA (Fig. 4A). Full-length RNA-Seq revealed marked differences in global gene expression levels, correlating with reporter activation either for both sensed target mRNAs, or specifically for IFI6 (Fig. 4B). While the former might comprise genes determining the state of cells in which CRISPR sensing works most efficiently, most of the specifically IFI6-coregulated genes were from the set of known ISGs. These results indicate a stochastic but correlated activation of ISGs across single cells at limiting IFNβ stimulation conditions.

**Figure 4.**
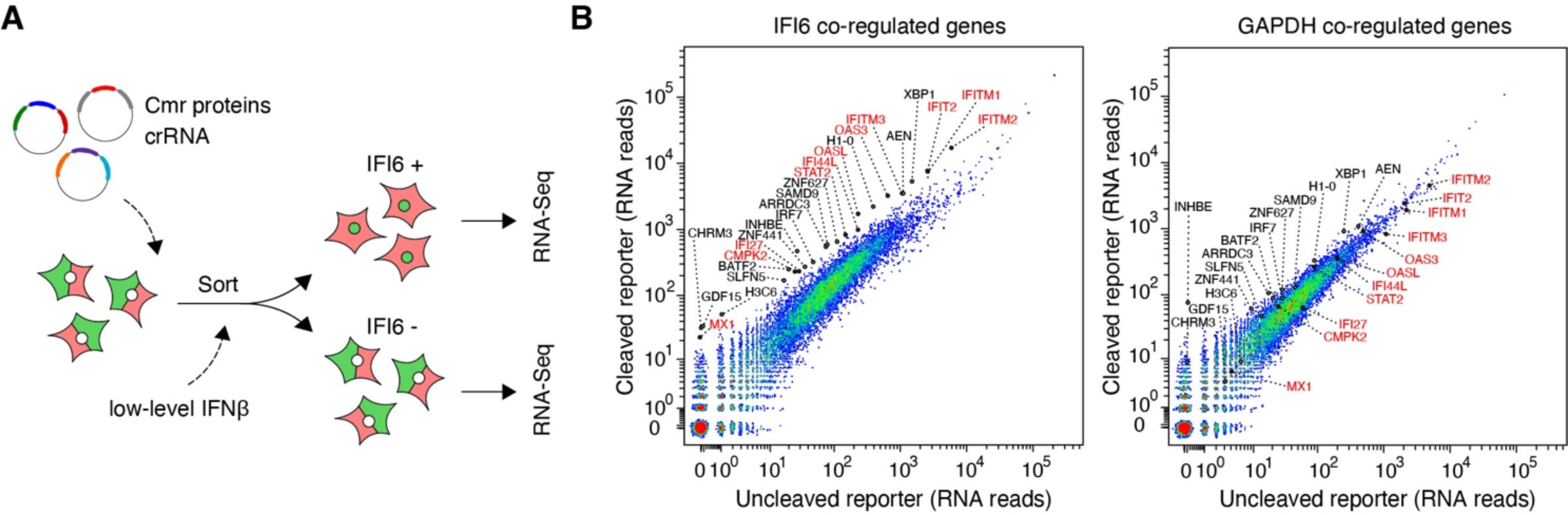
Detection of correlated stochastic gene expression in human cells using programmable CRISPR sensing. **(A)** Outline of detecting co-regulated gene expression by bulk transcriptomics after sorting on programmable mRNA sensing. **(B)** Stochastic co-expression analysis was performed by low-level stimulation of human cells with interferon and sorting based on ISG-or house-keeping control mRNA sensing. Enriched cells were analyzed by bulk transcriptomics. Highlighted are genes which are expressed higher in CRISPR-READ positive cells, indicating their co-regulation across single cells.

## DISCUSSION

In this study, we have successfully installed an orthogonal second messenger-mediated signaling pathway in human cells, which can be flexibly programmed to sense putatively any single-stranded RNA of known sequence in the cytosol. Unexpectedly, endogenous RNA sensing was only detectable when co-expressing four proteins of the *Thermus thermophilus* Cmr complex and immature crRNAs flanked by two repeats together with Cas6. The most likely explanation is that the maturation of crRNAs by Cas6 results in different RNA termini as compared to the mature crRNAs expressed from a U6 promoter, which in contrast could only sense overexpressed target RNAs.

It remains to be tested if this system can also sense nuclear, partially double-stranded, circular, short, non-coding, or highly modified RNA -In each of these cases, CRISPR-READ could provide a key resource to study previously hardly accessible areas of RNA biology in living cells. By optimizing the target specificity profile towards single-base discrimination, this system could be used to track cancer-associated somatic mutations or RNA deamination events in single living cells. Co-delivering a genome-or RNA editing system, CRISPR sensing could be used to enrich living cells with at least one edited allele, or lacking a non-edited allele.

We successfully performed genome-scale perturbation screening to identify genetic factors controlling a transcriptional program in cells. This screen will be easy to adapt to other transcriptional programs due to the flexible programming of CRISPR-mediated RNA sensing. Of note, such screens will not only provide a comprehensive picture of factors regulating transcription directly, but will also cover genes required for alternative splicing, nuclear export, and transcript stability.

Measuring correlated stochastic gene expression using CRISPR transcript sensing will help to understand the function of randomness in biology: are transcriptional bursts occurring at truly random moments like radioactive decay events, or are different fluctuating transcripts correlated due to a common control mechanism? Do inter-cellular differences in transcriptomic profiles serve a specific function, e.g., making the population of cells more resilient to stressors? Mapping out correlations between stochastic transcripts might reveal unexpected disease etiologies and novel drug targets.

The system we have engineered consists of seven proteins and one non-coding RNA. To realize its full potential, the system has to be delivered to primary cells or whole organisms using lentivirus, AAV, baculovirus, or large dsDNA integration. To reduce size, smaller orthologues could be screened for sensing ability, or ORFs could be linked by 2A peptides without individual promoters. A second color channel sensing another target RNA independently could be developed using components and second messengers from other species ^18^, which will enable to assess co-regulation of gene expression directly across single cells.

Time resolution of CRISPR sensing will be subject to engineering in two directions: a fast off-kinetics could be realized by fusing a proteasomal degradation domain to the fluorescent reporter, which could help following RNA dynamics at high time resolution; on the contrary, highest sensitivity will likely depend on inherent memory-like properties of CRISPR-READ. Lastly, an unexplored application of this system will be unlocked through delivering sensing crRNAs using a lentiviral library, rendering every cell a sensor for a different transcript. FACS sorting activated reporter cells and sequencing integrated crRNAs might elucidate transcriptome dynamics at high depth.

## Acknowledgements

We thank Jonathan Strecker and Thomas Ebert for continuous helpful discussions. J.L.S.-B. was supported by Deutsche Forschungsgemeinschaft (DFG) under Germany’s Excellence Strategy – EXC2151 – 390873048, Deutsche Forschungsgemeinschaft (DFG) SFB 1454 - project 432325352, and TRA Life & Health (University of Bonn). G.H. was supported by DFG grant: HA6805/6-1.

## Author contributions

J.L.S.-B. conceived of the study. K.B., G.H., and J.L.S.-B. designed the experiments. K.B., L.H., C.F., and N.S. collected the data. K.B., A.H., and J.L.S.-B. analyzed the data. K.B., A.H., G.H., and J.L.S.-B. wrote the manuscript with input from all coauthors.

## METHODS

### Mammalian cell culture

HEK 293T cells were cultivated in Dulbecco’s modified Eagle medium (DMEM) Glutamax media, additionally supplemented with 10% (v/v) fetal calf serum (FCS) and 10 μg/ml Ciprofloxacin in a 37°C incubator with 5% CO2.

### Golden Gate assembly

Sense and antisense oligos were annealed at a final concentration of 10 μM in T4 ligation buffer. If more than one insert was used, T4 polynucleotide kinase was added additionally during the annealing step to phosphorylate DNA ends for subsequent ligation. In a one-pot reaction, the backbone plasmid was digested with Esp3I and oligos were inserted and ligated using T7 ligase in StickTogether DNA Ligase Buffer (NEB). To ensure efficient ligation, DTT and BSA were added additionally at concentrations of 1 mM and 0.1 mg/ml, respectively. Sequence verification of transformants was performed by Tn5-mediated whole-plasmid tagmentation and MiSeq sequencing.

### KLD cloning

Plasmids were amplified with Q5 High-Fidelity 2X Master Mix (NEB) according to the manufacturer’s protocol, using a plasmid concentration of 0.4 ng/μl. The linearized plasmids were circularized using T4 PNK, T4 DNA Ligase, and DpnI in T4 Ligase buffer. Sequence verification of transformants was performed by Tn5-mediated whole-plasmid tagmentation and MiSeq sequencing.

### Generation of a stable reporter cell line

A stable reporter cell line was created using the piggyBac transposon system. HEK 293T cells were seeded at a density of 10^6^ cells/well in 6-well plates and transfected with 4 μg pPB_reporter_CalpL (Suppl. Table 1) and 1 μg pPiggyBacTransposase (Suppl. Table 1) per well using Lipofectamine 2000 (Invitrogen). After 48 hours, Blasticidin S selection was initiated at a concentration of 15 μg/ml. Cells were monitored until all untransfected control cells were dead.

### Live RNA sensing

Reporter cells were seeded at a density of 20,000 cells/well in 96-well plates. The following day, cells were transfected with a total of 100 ng of Cmr plasmids (20 ng Cmr2, 20 ng Cmr3, 30 ng Cmr4, 30 ng Cmr5 or 50 ng Cmr2_Cmr3_iRFP670 and 50 ng Cmr4_Cmr5_Cas6, Suppl. Table 1), 30 ng of crRNA expression plasmid, and 50 ng of target RNA expression plasmid, unless indicated otherwise, using 0.3 μl GeneJuice transfection reagent (Merck Millipore) per well. The expression of the reporter was induced by adding 1 μg/ml Doxycycline to the cells. Where indicated, IFNβ was added 2-3 hours post-transfection. After 48 hours, cells were harvested and analyzed via flow cytometry using a Miltenyi MACSQuant Analyzer equipped with a violet, cyan, and green laser.

### Flow cytometry analysis

Analysis of FCS files was performed using a custom open-source online application (https://jsb-lab.bio/xyplot/). Live cells were gated manually in forward- and side-scatter, followed by gating on cells expressing the reporter (GFP channel > 3,500). Depending on the experiment, positively transfected cells were additionally gated by iRFP670 expression, while gates were adjusted based on non-transfected control cells. Cells transfected with Cmr plasmids but no crRNA served as a negative control to distinguish FRET positive from FRET negative cells.

### Lentivirus-mediated cell line generation

sgRNAs were cloned into CROPseq-Guide-Puro (Addgene #86708). 10^6^ HEK 293T cells/well were transfected in 6-well plates with 1,760 ng lentiviral sgRNA plasmid along with lentiviral packaging plasmids pMD2.G (880 ng) and psPAX2 (1,320 ng) using Lipofectamine 2000. After 4-6 h, the medium was replaced. After 48 hours, the virus-containing supernatant was filtered through a 0.45 μm filter (Merck Millipore) and stored at -80°C. Reporter cells (8 × 10^6^ cells/well) were seeded in 6-well plates and transduced with 200 μl of lentivirus in the presence of 10 μg/ml polybrene (Merck Millipore). The following day, Puromycin selection was initiated at a concentration of 3 μg/ml (Cayman Chemicals). Cells were monitored until non-transduced control cells were dead.

### CRISPR knock out generation

An SpCas9 expression plasmid based on pcDNA3.1 was constructed (pCas9, Suppl. Table 1) and transfected transiently into sgRNA-expressing cells. In brief, 20,000 cells/well were plated in 96-well plates and transfected with 150 ng pCas9 plasmid per well, using GeneJuice transfection reagent (Merck Millipore). After recovery, cells were re-seeded and transfected with Cmr- and crRNA expression plasmids as detailed above. 48 hours post-transfection, cells were harvested, several replicate wells were pooled and sorted on FRET positive and FRET negative cells. Sorted cells were pelleted and lysed in 50 μl of direct lysis buffer ^30^ at 65°C for 10 minutes, and 95°C for 15 minutes. The targeted locus and/or lentiviral sgRNA was amplified and barcoded using two rounds of NEBNext PCR (NEB) (Suppl. Table 1) in two replicates per condition, and sequenced on an Illumina MiSeq.

### Genome-scale CRISPR screening using CRISPR RNA sensing as readout

For each of four screening replicates, 8 × 10^6^ reporter cells were transduced in 6-well format with 1 ml Human Brunello CRISPR knockout pooled library (Addgene #73178 ^31^) in the presence of 10 μg/ml polybrene (Merck Millipore). Starting after one day, cells were selected with 3 μg/ml Puromycin (Cayman Chemicals). Cells were split 1:3 when reaching confluency. After 4-7 days of selection, one million cells were lysed in 100 μl of direct lysis buffer at 65°C for 10 minutes, and 95°C for 15 minutes ^30^. Genome-integrated guide sequences were amplified and barcoded using two rounds of NEBNext PCR (NEB) and staggered guide-specific primers (Supplementary Table 1) in two replicates per library. After purification and Nanodrop-based quantification, libraries were sequenced on an Illumina NextSeq 2000 using a P2 100-cycle kit. Library members were counted using our open-source online application at www.jsb-lab.bio/LibCounter.htm.

### RNA-Seq

Cell pellets were resuspended in 350 μl RLT buffer (Qiagen) supplemented with 40 mM DTT. RNA was isolated using the NucleoSpin RNA Mini Kit (Marchery-Nagel) according to manufacturer’s protocol. Full-length RNA-Seq was performed using the SmartSeq2 protocol adapted for bulk RNA sequencing ^32^. Whole transcriptome amplified products (WTA) were tagmented using in-house produced Tn5 transposase followed by a barcoding PCR. After purification and quantification, libraries were sequenced on an Illumina NextSeq 2000 using a P2 100-cycle cassette.

### Imaging

All images were acquired using a Nikon Ti2 body equipped with a Yokogawa CSU-W1 spinning disc unit connected to Lumencor Celesta multimode lasers with wavelengths of 405 nm (nuclear staining), 477 nm (GFP), and 546 nm (mCherry). Emission filters used were Chroma ET450/50 (nuclear staining), Chroma ET525/50 (GFP), 572/28 BrightLine HC (mCherry). The exposure time was 90 ms for all channels. Objectives used were a Nikon 20x CFI P-Apo with a 1.5x tube lens inserted into the light path. A Hamamatsu Orca Flash4.0 LT+ camera was used in electronic shutter mode at full resolution (2048×2048).

## Data and material availability

NGS data will be available through SRA. Microscopy and FACS raw datasets are available upon request. All plasmid sequences and cloning strategies are provided in Suppl. Table 1. CRISPR-READ plasmids will be distributed through Addgene.org.

## Code availability

Custom analysis tools are available as open-source web applications at *www.jsb-lab.bio/xyplot/* and *www.jsb-lab.bio/LibCounter.htm*.

---

**Supplementary Figure 1.**
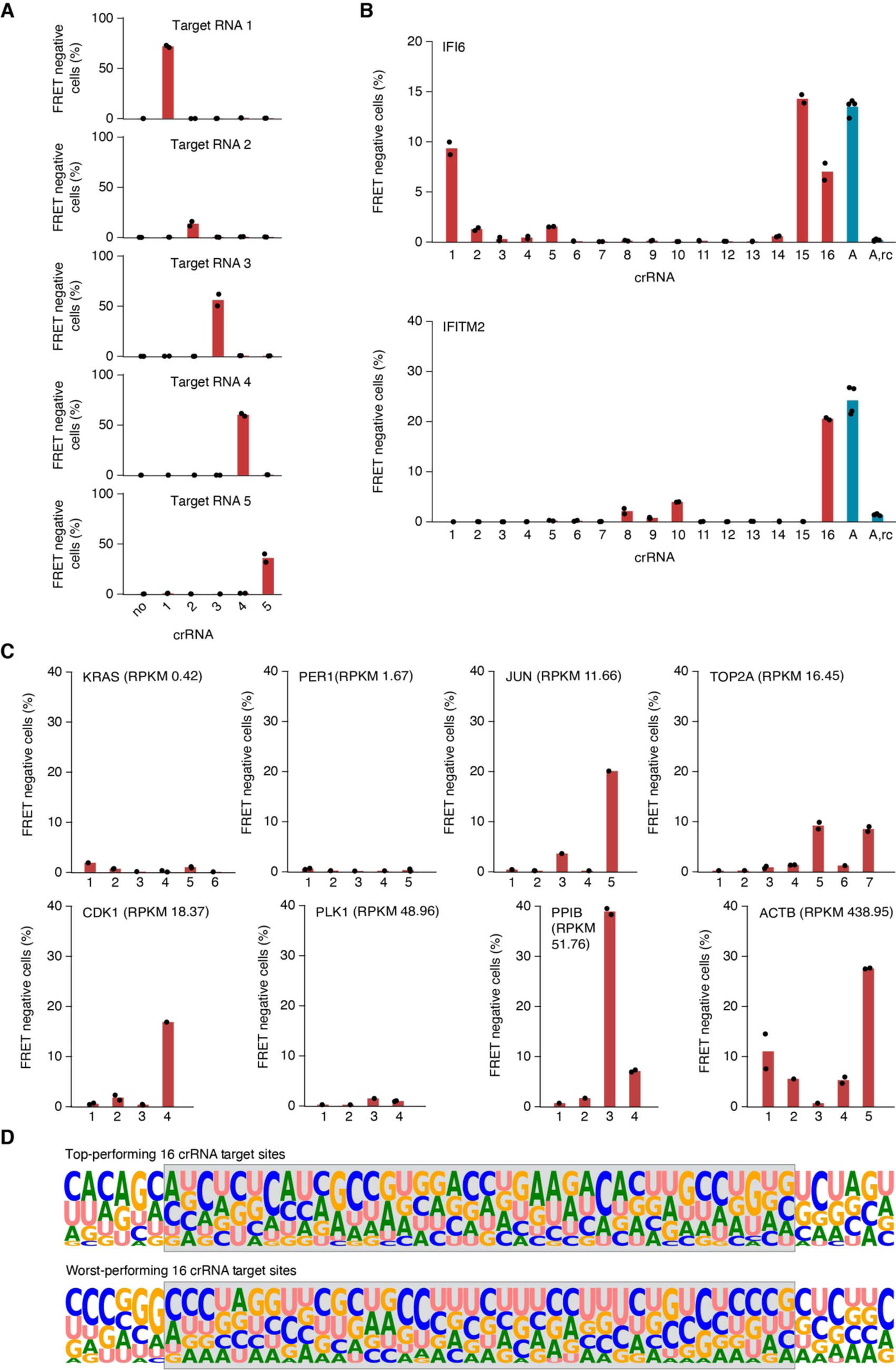
Tiling of additional human endogenous mRNAs. **(A)** Sequence-specific programmable RNA sensing upon expression of matching and mismatching pairs of double repeat-flanked crRNAs and target RNAs under a polymerase III promoter. Shown are fractions of FRET-negative cells measured by FACS. **(B)** Sensing efficiency of single crRNAs tiling the IFI6- or IFITM2 mature mRNA sequence (red bars) or repeat-spacer arrays targeting the same mRNA or its reverse complement (blue bars). **(C)** Sensing efficiency of single crRNAs tiling the indicated genes with indicated expression levels HEK 293T cells measured by full-length RNA-Seq. **(D)** Sequence logos of the 16 best or worst performing CRISPR-READ target sequences out of 81 sequences tested. The crRNA binding site is highlighted in grey.

